# Neuronal acid-sensing ion channel 1a regulates neuron-to-glioma synapses

**DOI:** 10.1101/2023.08.31.555794

**Authors:** Gyeongah Park, Zhen Jin, Qian Ge, Yuan Pan, Jianyang Du

## Abstract

Neuronal activity promotes high-grade glioma progression via secreted proteins and neuron-to-glioma synapses, and glioma cells boost neuronal activity to further reinforce the malignant cycle. Whereas strong evidence supports that the activity of neuron-to-glioma synapses accelerates tumor progression, the molecular mechanisms that modulate the formation and function of neuron-to-glioma synapses remain largely unknown. Our recent findings suggest that a proton (H^+^) signaling pathway actively mediates neuron-to-glioma synaptic communications by activating neuronal acid-sensing ion channel 1a (Asic1a), a predominant H^+^ receptor in the central nervous system (CNS). Supporting this idea, our preliminary data revealed that local acid puff on neurons in high-grade glioma-bearing brain slices induces postsynaptic currents of glioma cells. Stimulating Asic1a knockout (Asic1a^-/-^) neurons results in lower AMPA receptor-dependent excitatory postsynaptic currents (EPSCs) in glioma cells than stimulating wild-type (WT) neurons. Moreover, glioma-bearing Asic1a^-/-^ mice exhibited reduced tumor size and survived longer than the glioma-bearing WT mice. Finally, pharmacologically targeting brain Asic1a inhibited high-grade glioma progression. In conclusion, our findings suggest that the neuronal H^+^-Asic1a axis plays a key role in regulating the neuron-glioma synapse. The outcomes of this study will greatly expand our understanding of how this deadly tumor integrates into the neuronal microenvironment.

## INTRODUCTION

High-grade gliomas are among the most lethal malignancies of the central nervous system (CNS), infiltrating and disrupting brain structure and function. Despite intensive research efforts, high-grade glioma survival has not changed significantly over the past two decades, indicating an urgent need to develop novel therapeutic strategies. Recent research indicates that neuronal activity may be a potential therapeutic target for high-grade gliomas (Johung and Monje, 2017).

As the field of cancer neuroscience grows, it has become increasingly clear that neurons influence glioma pathophysiology. For example, neuronal activity is required for tumor initiation and maintenance in low-grade gliomas (Pan et al., 2021); and other studies uncovered that neuronal activity drives high-grade glioma growth and migration via bona fide neuron-to-glioma synapses (Venkataramani et al., 2019; Venkatesh et al., 2019; Taylor et al., 2021; Barron et al., 2022; Venkataramani et al., 2022; Huang-Hobbs et al., 2023). The functional neuron-to-glioma synapses trigger the downstream calcium (Ca^2+^) signaling in glioma cells through activating glioma AMPA receptors (AMPARs) (Ishiuchi et al., 2002; Ishiuchi et al., 2007). In addition, glioma cells produce factors (*e.g.* glypican-3) that increase synaptogenesis in the tumor microenvironment, leading to neuronal hyperexcitability that further enhances the activity-dependent high-grade glioma growth (Buckingham et al., 2011; Campbell et al., 2012; John Lin et al., 2017; Venkatesh et al., 2019; Hatcher et al., 2020; Yu et al., 2020). Whereas the importance of neuron-glioma interaction in glioma pathophysiology has been established, many regulators of this interaction remain to be discovered. Uncovering critical regulators of neuron-glioma synaptic communications will ultimately guide the development of strategies against this deadly cancer.

Recent studies have demonstrated that extracellular tumor acidification is a common feature of glioma growth (Honasoge and Sontheimer, 2013; Rohani et al., 2019). This acidification is evident in the glioma cells of fast proliferation, which results in the production of significant amounts of metabolic waste products including lactate and carbon dioxide by consuming nutrients and oxygen from the tissues around them (DeBerardinis et al., 2007; Lugano et al., 2020; Tanaka et al., 2021). This acidification has a range of pathophysiological consequences, including disrupting normal cellular metabolism, altering the function of various proteins and enzymes, and promoting the growth and survival of glioma cells (Pienkowski et al., 2021; Alzial et al., 2022; He et al., 2022). Consolidated data from multiple investigations have revealed that tumor masses can have varied pH levels ranging from 7.4-6.2 units and can decrease to 5.5-3.4 units or even below (Webb et al., 2011; Kato et al., 2013; Chen et al., 2014).

The acidic local environment can influence surrounding neurons by activating pH-sensitive membrane proteins including receptors (Chesler, 2003; Sinning and Hubner, 2013). One of the dominant H^+^ receptors on neurons is ASICs. ASICs are a group of non-selective cation channels that are activated by acidic pH levels (increased H^+^ concentration). The ASIC family contains six identified genes (*ASICs 1a, 1b, 2a, 2b, 3, and 4*) (Waldmann et al., 1997; Waldmann and Lazdunski, 1998). In the brain, Asic1a and Asic2 are the two major subunits that assemble as homo- or hetero-trimers to form acid-activated cation channels (Wemmie et al., 2013). The activation of neuronal ASICs by extracellular acidic pH leads to sodium (Na^+^) and/or Ca^2+^ influx, resulting in neurotransmitter release and the initiation of signaling pathways (Wemmie et al., 2013; Du et al., 2014). ASICs have been implicated in various neurological disorders such as ischemic injury, chronic pain, and psychiatric disorders (Xiong et al., 2004; Mazzuca et al., 2007; Coryell et al., 2009; Ziemann et al., 2009).

Interestingly, some studies have found that Asic1a signaling can enhance the differentiation of neural stem cells into astrocytes, which could have implications for the role of Asic1a in glioma progression (Barkho et al., 2006; Luo et al., 2017; Ha et al., 2019; Magnusson et al., 2020; Nistor-Cseppento et al., 2022). Furthermore, Asic1a signaling can promote glioma proliferation via extracellular acidifications, potentially through its regulation of intracellular Ca^2+^ levels (Takayasu et al., 2020; Elias et al., 2023). These studies imply that glioma-initiating cells may be particularly susceptible to changes in ion homeostasis, such as intracellular Ca^2+^ and potassium (K^+^) levels. Although glioma-expressed Asic1a might regulate tumor progression, controversial pro- and anti-cancer effects were reported (Honasoge and Sontheimer, 2013; King et al., 2021; Sheng et al., 2021). Alternative mechanisms are required to explain how glioma acidification governs its pathophysiology, and further research is needed to better understand the complicated relationships between Asic1a and the microenvironment of glioma, in which neuron-glioma synaptic communication appears to be key.

Collectively, a greater understanding of the mechanisms by which the H^+^-Asic1a axis regulates neuron-glioma synaptic communication has the potential to provide new insights into glioma pathophysiology as well as uncover new targets for therapeutic intervention.

## MATERIALS AND METHODS

### Mice and ethic statement

All mice were maintained on a congenic C57BL/6 background. Asic1a^-/-^ mice were described previously (Wemmie et al., 2002). Experimental groups were matched for age (8-10 weeks old). Mice were housed on a standard 12-hour light-dark cycle and received standard chow (LM-485; Teklab, Madison, WI, USA) and water ad libitum. The University of Tennessee Health Science Center Animal Care and Use Committee approved all procedures. All efforts were made to minimize suffering.

### GL261 glioma cell culture

The GL261 murine glioma cell line (NCI Tumor Repository, Frederick, MD) was grown in Dulbecco’s modified Eagle’s medium (DMEM) with 10% fetal bovine serum (FBS), 100 U/ml penicillin, and 100 μg/ml streptomycin and grown at 37°C in a 5% CO_2_ and humidified atmosphere, all obtained from Thermo Fisher Scientific (Waltham, MA, USA).

### Generating a GFP-labeled GL261 (GL261-GFP) cell line

GL261 cells were transfected with a green fluorescent protein (CopGFP) lentiviral particle (sc-108,084; Santa Cruz Biotechnology (Santa Cruz, CA, USA) according to the manufacturer’s protocol. We infected the GL261 cells by adding the lentivirus to the glioma cell culture medium with polybrene (optional, 10 μg/ml working concentration, Sigma) to increase the efficiency of infection. The cell culture medium was replaced with a fresh medium 24 hours after the virus infection. The virus-infected GL261 cells were trypsinized with 0.25% trypsin (Sigma-Aldrich) 72 hours after infection. The viable cells with bright fluorescence were sorted out using a FACSAria cell sorter (BD Biosciences) in the UTHSC flow cytometry and cell sorting laboratory. Briefly, to set the gate of the background signal, control GL261 cells without GFP expression were placed onto the fluorescence-activated cell sorting. GFP-expressing cells were sorted based on the intensity of their GFP signals. The sorting gate was configured to gather the 10% of the cell population that fluoresced the most with GFP. The sorted cells were planted in DMEM on a 24-well plate for amplification and maintenance.

### In vivo GL261-GFP cell implantation in mice

For cell implantation, cells were trypsinized with 0.05% trypsin-EDTA washed and resuspended in phosphate-buffered saline (PBS) at a final concentration of 1×10^4^ cells/μl. All animals were anesthetized with 2% isoflurane mixed with oxygen and were positioned in a stereotaxic (RWD life science) head frame and anesthesia mask. Following sanitation of the surgical area, a single skin cut of about 0.5 cm was performed by using scissors, extending from the bregma to the lambda suture. We injected GL261 cells, which exhibit high-grade glioma features, into the mouse amygdala. A 2.0 mm hand drill was used to drill a hole in the skull at the injection site, relative to bregma: mediolateral (ML) ±3.5 mm; anteroposterior (AP) -1.2 mm; dorsoventral (DV) -4.3 mm. Using a 10 μl Hamilton syringe with a syringe pump (Legato-130, KD scientific), a 2 μl GL261-GFP cell suspension was stereotactically injected into the brain. The injection took about 10 minutes to complete, including 5 minutes of holding time and 1 minute to remove the needle. The mice were placed on a heating plate after surgery until they were awakened. To release the pain, buprenorphine (3 mg/g body weight) was administered, and their postoperative condition was monitored daily. To ensure control over the time difference between injections, ensuring that injection timing is constant throughout all groups, a mouse from each group was injected before another mouse from the same group during the process.

### Animal survival study

We injected GL261 or GL261-GFP cells (1×10^4^ cells/ml in PBS) into the amygdala of WT, Asic1a^-/-^, Asic1a^f/f^, and Asic1a^Vglut1-cKO^ mice, respectively (as depicted along the stereotactic axis above). Following the surgical procedure, the mice’s body weight was monitored every 3 days. In the event of any signs of distress or neurological symptoms, and if any mice met the humane endpoint criteria, they were then humanely euthanized (Ullman-Cullere and Foltz, 1999; Stokes, 2002). To assess survival rates, Kaplan-Meier survival analysis with log-rank testing was conducted for statistical evaluation (Dinse and Lagakos, 1982; Goel et al., 2010).

### Brain slices preparation for patch-clamp

In cutting solution for brain preparation contained (in mM): 205 sucrose, 5 KCl, 1.25 NaH_2_PO_4_, 5 MgSO_4_, 26 NaHCO_3_, 1 CaCl_2_, and 25 glucose. The artificial cerebrospinal fluid solution (ACSF) for whole-cell recording contains (mM): 115 NaCl, 2.5 KCl, 2 CaCl_2_, 1 MgCl_2_, 1.25 NaH_2_PO_4_, 17 glucose, 25 NaHCO_3_ bubbled with 95% O_2_/5% CO_2_, pH 7.4 at 23 ± 3°C. The pipette solution for whole-cell patch-clamp recordings contains (in mM): 145 Cs-methanesulfonate (CsSO_3_CH_3_), 8 NaCl, 1 MgCl_2,_ 0.38 CaCl_2_, 1 Cs-EGTA, and 10 HEPES (mOsm 290-310, adjusted to pH 7.25 with CsOH). The pipette resistance was 8-10 M ohm. The brain was extracted from the skull and coronal sections (300 μm thick) containing GL261-GFP in the amygdala were sliced using a vibratome (Leica VT1000s). Slices were bubbled with 95% O_2_/5% CO_2_ for one hour at ambient temperature (22 ± 3°C). To perform the whole-cell patch-clamp recording, the slices were placed in a submerged recording chamber and perfused with ACSF at a rate of 2 ml/min. In some experiments, for injections of acidic ACSF into the slice, acidic pH solutions were prepared as described previously (Du et al., 2014). In brief, the pH was buffered by 10 mM HEPES in solutions with a pH > 6.0, and by 10 mM MES when the pH was less than 6.0. Three to five-second puffs of acidic solution were delivered by a patch-clamp pipette attached to a microINJECTOR system (Tritech Research).

### Patch-Clamp Recording

We collect electrophysiological data from brain slices three to seven days after implanting GL261-GFP cells. The glioma cell and neuronal activities in the amygdala brain slices were recorded through an Axon Instruments Multiclamp 700B amplifier, sampled at 10 kHz, and digitized with a Digidata 1440A (Molecular Device software). The cell morphology was identified under an Olympus BX51 fluorescence microscope. A bipolar electrode was placed at cortical inputs to evoke AMPA-dependent EPSCs. CNQX (100 µM, BioGems) was used to inhibit AMPA currents. Voltage-dependent K^+^ currents were measured from -70 to +90 mV with a 20-pA step in each recording and 4-AP (200 µM, Sigma-Aldrich) was used to block K^+^ currents. mEPSCs were recorded with holding potential at -80 mV, in the presence of 100 µM picrotoxin (Acros organics) to block GABA_A_ receptors. The puff of acid was placed on cortical inputs or neuronal soma to activate neuronal activity. PcTX1 (200 ng/ml, Peptide Institute, Inc) was used to block Asic1a-dependent currents. In most cases, we analyzed current amplitude (pA) with Clampfit 11.2. The AMPA-and K^+^-dependent currents were analyzed and displayed by current density (pA/pF).

### Western blot Analysis

Fourteen days after the GL261-GFP cell injection, we examined western blot and immunostaining. GL261 cells were lysated in a RIPA buffer (50 mM TrisHCl, 150 mM NaCl, 1% Triton, 0.1% SDS, 1% deoxycholate, 2 mM EDTA, pH 7.5) added with a protease and phosphatase inhibitor cocktail (Thermo Fisher Scientific, IL, USA). Protein samples were separated by 10% SDS-polyacrylamide gel electrophoresis (20 µg/lane). The blot has been cut probing different regions of the same blot with the following primary antibodies: rabbit anti-Asic1a (1:1000, Invitrogen), rabbit anti-GluA1 (1:1000, Cell Signaling), rabbit anti-GluA4 (1:1000, Cell Signaling), rabbit anti-Neuroligin-3 (1:1000, Thermo Fisher Scientific), rabbit anti-COX-2 (1:1000, Thermo Fisher Scientific), mouse anti-MGMT (1:1000, Invitrogen), and rabbit anti-ꞵ-actin (1:2000, Cell Signaling). Detection was performed through the chemiluminescence assay with an ECL plus kit (Thermo Scientific, IL, USA). Densitometric detection was carried out with the LI-COR Odyssey imager and Image Studio software (version 5.2, LI-COR Biosciences, USA).

### Tissue collection and immunohistochemistry

Mice were anesthetized with 2% isoflurane followed by transcardial perfused with 4% paraformaldehyde in 0.1 M PBS. After six hours of fixation, mouse brains were immersed in 15%, 20%, and 30% sucrose at 4°C until they sank. Brains were sliced into 30 μm thick coronal sections and stained with cresyl violet (Nissl). Brain sections were incubated with rabbit anti-Neuroligin-3 (1:250, Thermo Fisher Scientific) overnight at 4°C. After washing with PBS, sections were incubated for one hour at room temperature with a secondary antibody. Sections were moved into gelatin-coated slides (Thermo Fisher Scientific) and were mounted onto cover slides (Electron Microscopy Sciences). Sections were visualized (10x and 20x magnification) under a Leica microscope (DMRXA2, Leica Microsystems) equipped with a digital camera and imaging software. Digital images were captured and saved. Immunohistochemistry analysis was performed using ImageJ software (NIH).

For Immunofluorescence, brain sections were incubated with primary antibodies overnight at 4°C. After washing with 0.1 M PBS, sections were incubated with Alexa Fluor 488-and 594-conjugated donkey secondary antibodies from Invitrogen. Prolong Gold antifade reagent with DAPI (Invitrogen) was used to label the nuclei. Fluorescence images of the sections were captured using a Leica microscope and saved. For each region, the ratio of fluorescence intensities was measured and calculated manually using ImageJ.

### Statistical analysis

Statistical data were analyzed using GraphPad Prism 8. There were no inclusion criteria defined, but membrane resistance and capacitance were measured. Data are expressed as the mean ± SEM of at least three independent experiments unless otherwise indicated. Two-way ANOVA and Tukey’s test were used for comparisons among multiple groups, and two-tailed Student’s t-test for used for comparisons between two groups. A statistically significant result was defined as a p-value of 0.05.

## RESULTS

### *Asic1a* deletion in the mouse brain reduces tumor growth and prolongs mouse survival

To evaluate the impact of brain *Asic1a* gene knockout on glioma growth in an immunocompetent context, we engineered the murine GL261 cells to express GFP (GL261-GFP). The GL261-GFP cells stably expressed GFP through fluorescence-activated cell sorting, even after 2 months of growth (**Fig. 1A**). Five to seven days after implanting GL261-GFP cells into the brain (amygdala) of C57BL/6J mice, all mice developed gliomas (> 2 mm in diameter).

**Figure 1.**
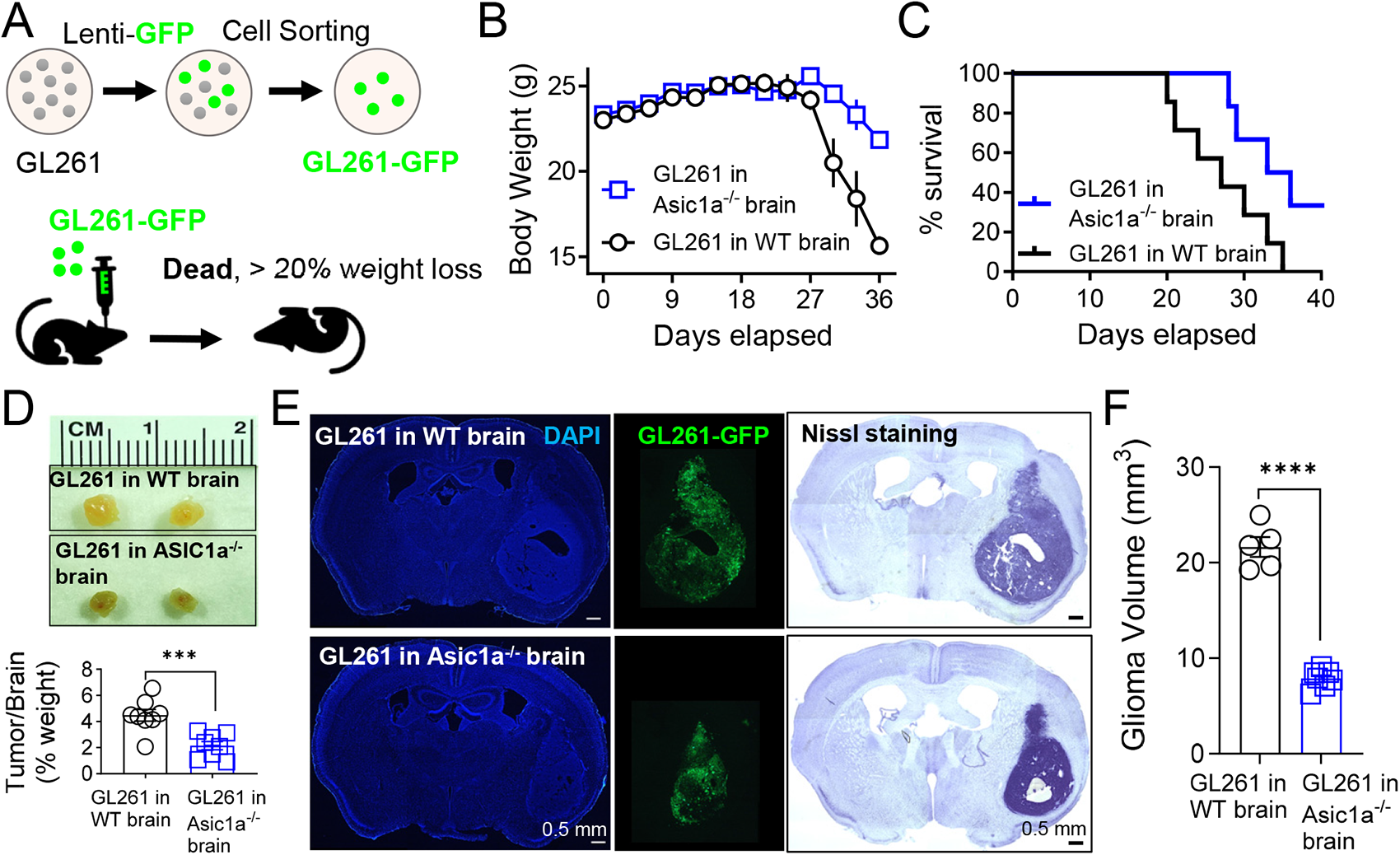
Brain Asic1a deletion reduces glioma growth and increases survival rate in glioma-bearing mice. **(A)** The schematic shows the experimental approach for creating a GFP-labeled GL261 glioma cell line. **(B)** Time-dependent body weight change in the glioma-bearing WT (black, *n* = 14) and Asic1a^-/-^ mice (blue, *n* = 16). **(C)** Time-dependent survival measurement in the glioma-bearing WT (black, *n* = 14) and Asic1a^-/-^ mice (blue, *n* = 16). **(D)** The glioma samples were collected from GL261-GFP-bearing WT mice and Asic1a^-/-^ mice, respectively. Tumor/brain weight comparison of G261-GFP in WT (black, *n* = 8) and in Asic1a^-/-^ mice (blue, *n* = 8). *** p = 0.0008 *(t = 4.239, df = 14)*. **(E)** The representative immunostaining images of DAPI (left), GFP-tagging G261 (middle), and Nissl stain (right) in G261-GFP-bearing WT (up) and Asic1a^-/-^ (down) brain slices at 14 days. **(F)** Glioma volume between G261-GFP in WT (black, *n* = 5) and in Asic1a^-/-^ mice (blue, *n* = 7), ****p < 0.0001 *(t = 14.25, df = 10)*. All statistics were analyzed by a two-tailed unpaired Student’s t-test. Data are mean ± SEM.

Following the implantation of GL261-GFP cells, several approaches were employed to comprehensively evaluate the influence of *Asic1a g*ene deletion. First, we engaged in electrophysiological measurements at the early stages, specifically spanning from 3 to 7 days post-injection (see Figs. 3 to 6). Subsequently, tissue samples were collected on day 14, allowing for a more in-depth analysis of the evolving dynamics within the brain tissue. For mice survival studies, we monitored the body weight and overall survival of the experimental subjects over a span of 36 days, as depicted in **Fig. 1B, C**. Initial observations revealed that there were no discernible differences between the two experimental groups in terms of body weight at the 14-day mark (**Fig. 1B, C**). This suggests that *Asic1a* deficiency did not manifest any immediate overt effects on these parameters during the initial stages of tumor development. However, a notable trend emerged in the subsequent days, showcased in **Fig. 1B, C**, wherein GL261-GFP implanted in the brains of WT mice experienced a significant reduction in body weight between days 30 and 36 compared to their GL261-GFP in *Asic1a* ^-/-^ counterparts (**Fig. 1B**). Of particular significance was the disparity in survival rates between the two experimental groups, as highlighted in **Fig. 1C**. It was evident that the absence of Asic1a in the brain led to a substantial increase in the overall survival with GL261-GFP implantation, suggesting a potential protective effect of neuronal *Asic1a* gene deletion in the context of tumor progression.

To further investigate tumor features, we dissected the tumors from the postmortem brains. The weight of the tumors is significantly lower in the *Asic1a* ^-/-^ groups versus the WT brain groups (**Fig. 1D**). We then employed Nissl staining to visualize glioma cell distribution and density. Our observations demonstrated a noticeable reduction in tumor volume within the brains of GL261-Asic1a^-/-^ mice compared to those in GL261-WT mice (**Fig. 1E, F**). These data demonstrate that *Asic1a* deletion grants a survival advantage to mice harboring high-grade gliomas.

To unravel the cell-type-specific mechanism underpinning how Asic1a modulates glioma progression, we employed a targeted approach involving the generation of Asic1a conditional knockout mice. This was achieved through a crossbreeding strategy that combined Asic1a^f/f^ mice with the Vglut1-IRES2-Cre-D transgenic mice, facilitating the elimination of Asic1a specifically in excitatory neurons (referred to as Asic1a^Vglut1-cKO^). Subsequently, GL261-GFP cells were implanted into the brains of Asic1a^Vglut1-cKO^ mice. Our investigations unveiled a reduced tumor size, as evidenced by Nissl staining, in the brains of Asic1a^Vglut1-cKO^ mice (**Fig. 2A**). Consistently, there was a ∼ 46% reduction in tumor volume within the brains of GL261-Asic1a^Vglut1-cKO^ mice (42.85 ± 1.20 mm^3^) compared to their GL261-Asic1a^f/f^ controls (19.59 ± 1.37 mm^3^) (**Fig. 2B, C**). Notably, the tumor-to-brain weight ratio pointed towards a ∼ 26% reduction in tumor growth within the Asic1a^Vglut1-cKO^ brains (WT: 2.72 ± 0.19 % versus Asic1a^Vglut1-cKO^: 2.13 ± 0.04 %, **Fig. 2F**).

**Figure 2.**
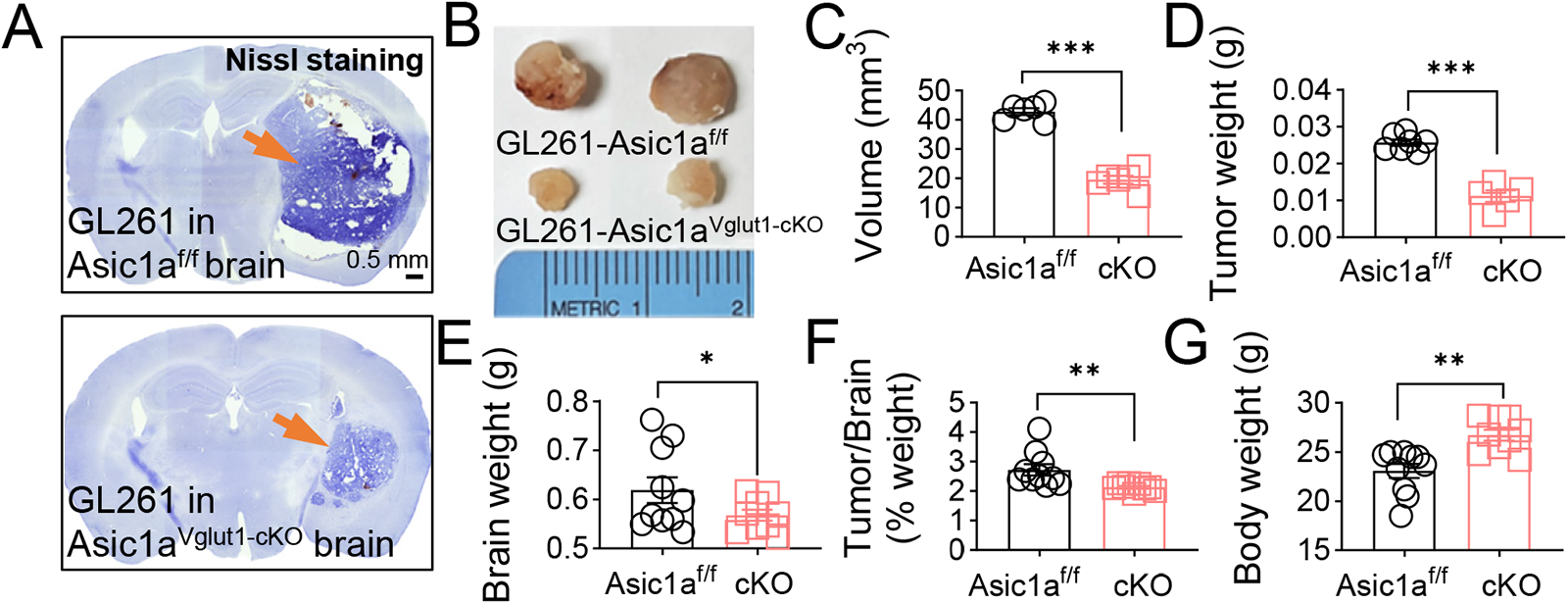
Excitatory neuronal Asic1a deletion reduces glioma growth. **(A)** The representative Nissl staining in GL261-GFP bearing ASIC1a^f/f^ (up) and Asic1a^Vglut1-cKO^ (down) brain slices. The arrows indicated the glioma in the amygdala slices. **(B)** The glioma samples were collected from GL261-GFP-bearing WT mice and Asic1a^Vglut1-cKO^ mice, respectively. **(C-G)** The comparison of tumor volume **(C)**, **** p < 0.0001 (*t=12.78, df=10)*; tumor weight **(D)**, **** p < 0.0001 *(t=8.978, df=10)*; brain weight **(E)**, ns, non-significant, p = 0.1076 *(t=1.698, df=17)*; Tumor/brain weight **(F)**, ** p = 0.0097 *(t = 2.911, df = 17)*; and body weight **(G)**, **p = 0.001 *(t = 3.946, df = 17)* from GL261 bearing Asic1a^f/f^ (black, *n* = 6-10) and Asic1a^Vglut1-cKO^ mice (red, *n* = 6-9). All statistics were analyzed by a two-tailed unpaired Student’s t-test. Data are mean ± SEM.

These outcomes closely paralleled those observed in the GL261-Asic1a^-/-^ mice (**Fig. 1**), suggesting a substantial impact of brain Asic1a on high-grade glioma progression, primarily through interactions between excitatory neurons and glioma cells. An intriguing observation emerged in the realm of physiological impacts, wherein the body weight of GL261-Asic1a^Vglut1-cKO^ mice was increased in comparison to their GL261-Asic1a^f/f^ counterparts (**Fig. 2G**). This finding suggests that the deletion of *Asic1a* specifically in excitatory neurons might confer a potential rescue effect on the body weight loss typically induced by glioma, thus implying an avenue of promise for clinical therapeutic interventions.

### *Asic1a* deletion in mouse brain reduces EPSCs in the implanted glioma cells

Numerous investigations have underscored the pivotal role of glutamate signaling, particularly involving the AMPA receptor (AMPAR), in shaping the intricate interplay between neurons and glioma cells (Venkataramani et al., 2019; Venkatesh et al., 2019; Taylor et al., 2021; Barron et al., 2022; Venkataramani et al., 2022; Huang-Hobbs et al., 2023). In light of this, our study aimed to delve into the role of glioma AMPARs in regulating the neuron-glioma crosstalk.

Building upon these observations, we formulated a hypothesis that the diminished connectivity between neurons and glioma cells in the context of *Asic1a* gene knockout would manifest as a reduction in AMPA-mediated glioma-EPSCs in glioma-Asic1a^-/-^ mice. The GL261-GFP cells were implanted into the amygdala for 3-7 days until patch-clamp recording (**Fig. 3A**). The co-localization of GL261-GFP and PSD-95 expression in the representative immunostaining image revealed an intergradation of glioma and neurons (**Fig. 3A**). To rigorously test this proposition, we simulated amygdala neurons to elicit and record EPSCs in the implanted visible GL261-GFP cells within the amygdala (**Fig. 3B**). Intriguingly, our investigations into the evoked EPSCs unveiled a significant role played by Asic1a in modulating the connection between glioma cells and neurons. We observed a discernible reduction in the density of EPSCs in the glioma implanted in Asic1a^-/-^ mice compared to their glioma-WT mice controls (**Fig. 3C, D**). The specificity of this phenomenon was validated through the blockade of evoked currents by CNQX, a pharmacological agent targeting AMPA receptors, thus confirming the AMPA-mediated nature of these currents (**Fig. 3E**). Conversely, no significant differences emerged in the paired-pulse ratio (PPR) between glioma-WT and glioma-Asic1a^-/-^ mice (**Fig. 3F, G**), suggesting that the alterations in the glioma-EPSCs may not have been caused by a change in the vesicle release patterns in the Asic1a^-/-^ brains. The conductance of the GL261-GFP cells remained unaltered when implanted in Asic1a^-/-^ brains versus in WT brains (**Fig. 3H**), demonstrating that Asic1a deletion in the brain has no effect on modifying glioma cell size, one of the markers of cellular differentiation (Chen et al., 2020).

**Figure 3.**
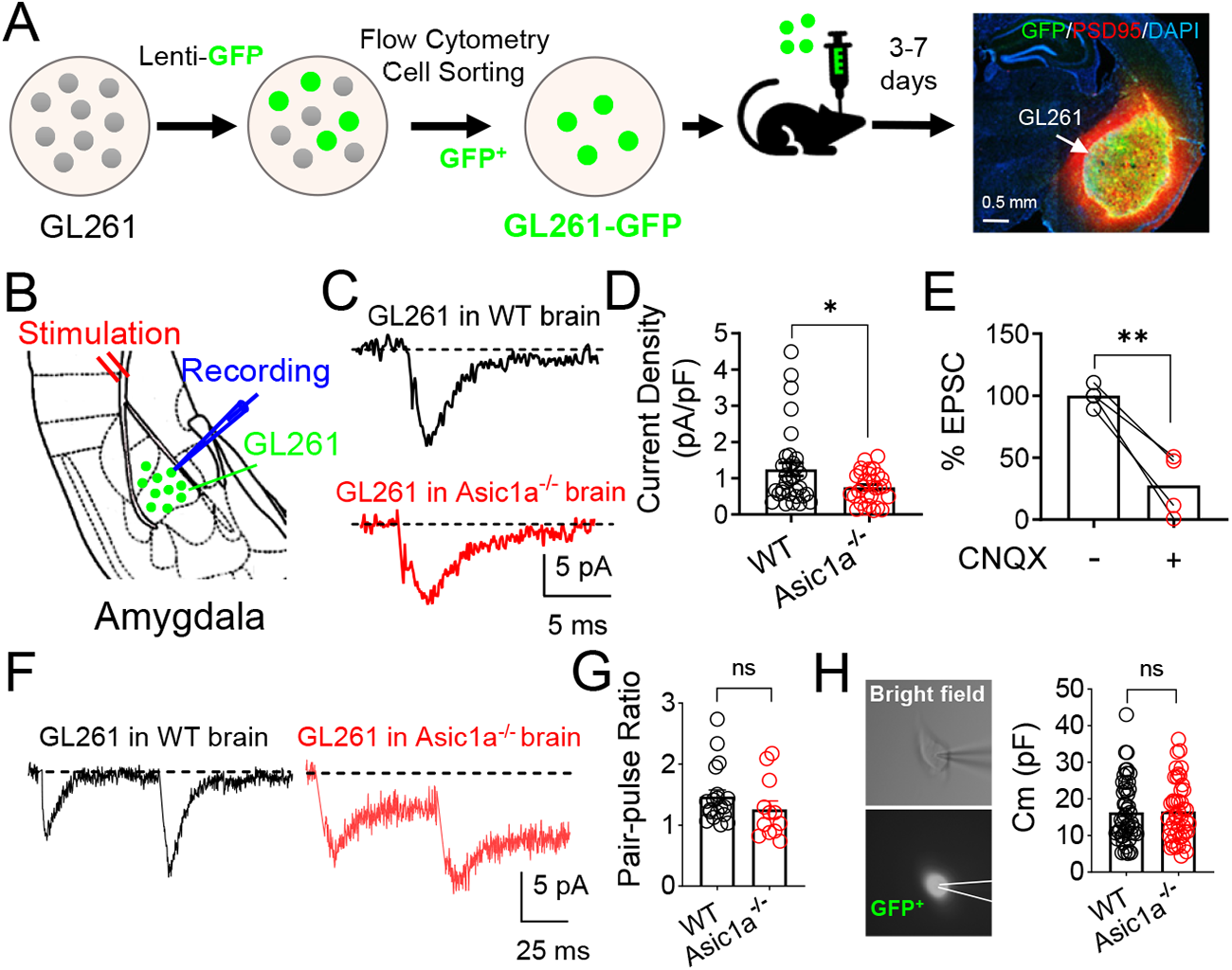
Brain Asic1a deletion attenuates glioma-EPSCs in the brain. **(A)** The schematic shows the creation of GL261-GFP cells and visual identification in the amygdala by immunofluorescence. GFP (green, GL261 cell), PSD 95 (red), and DAPI (blue). **(B)** The schematic of EPSC recording in GL261 by stimulating amygdala cortical input. **(C)** The representative trace of evoked GL261-EPSCs in WT (black) and Asic1a^-/-^ (red) amygdala slices. **(D)** The EPSC density (pA/pF) in GL261-GFP in WT (*n* = 32 cells in 8 mice) and Asic1a^-/-^ (*n* = 29 cells in 8 mice) amygdala slices. * p = 0.022 *(t = 2.359, df = 60)*. **(E)** The evoked EPSCs were inhibited by 100 µM CNQX (*n* = 4 cells in 4 mice). ** p = 0.006 *(t = 6.815, df = 3)*. **(F)** The representative paired-pulse ratio (PPR) in GL261 in WT (black) and ASIC1a^-/-^ (red) amygdala slices. **(G)** The summarized PPR in G261 in WT (*n* = 19 cells in 8 mice) and ASIC1a^-/-^ mice (*n* = 12 cells in 8 mice). non-significant, p = 0.2328 *(t=1.219, df=29)*; **(H)** Left, representative bright field (up) and GFP (down) images of GFP-tagging cells in the brain. Right, the cell conductance measurement of the glioma. ns., non-significant, p = 0.8943 *(t=0.1332, df=94)*; All statistics were analyzed by a two-tailed unpaired Student’s t-test. Data are mean ± SEM.

To deepen our understanding of the functional implications within the neuron-glioma interaction context, we proceeded to investigate miniature EPSCs (mEPSCs) in each experimental group (**Fig. 4A**). Interestingly, our exploration of mEPSCs in GL261-GFP cells revealed decreased frequency and amplitude in the GL261 cells implanted in Asic1a^-/-^ brains compared to the WT counterparts (**Fig. 4B-D**). To further identify whether the recorded EPSCs are AMPAR dependent, we evaluated glutamate receptor expression in GL261 cells and cultured SY5Y neuronal cells. Our data indicated that GluA1 and GluA4 subunits were present in GL261 cells (**Fig. 4E, F**). These findings suggest that sufficient and functional AMPARs may participate in the neuronal activity-dependent EPSCs in GL261 cells.

**Figure 4.**
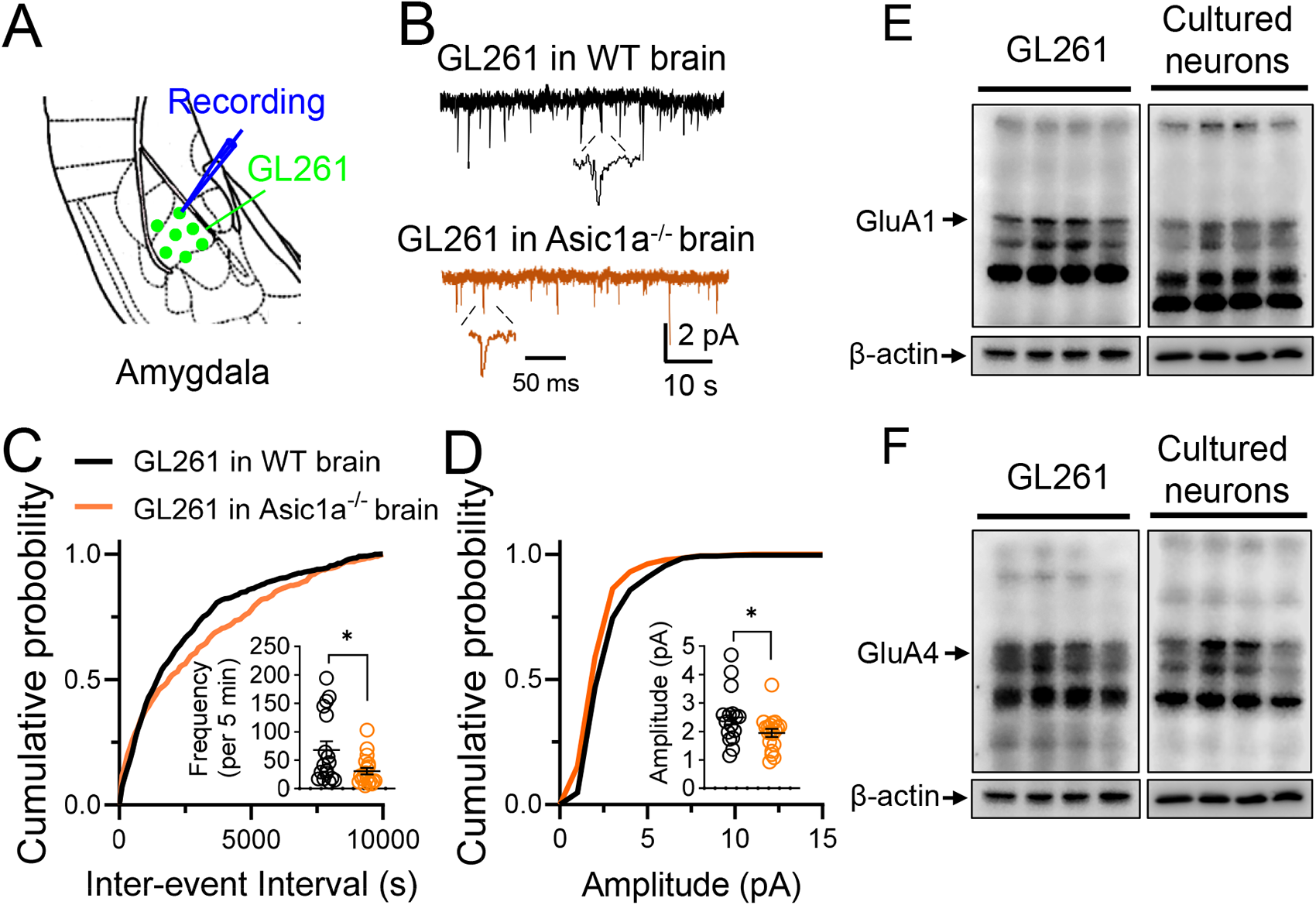
Brain Asic1a deletion attenuates glioma-mEPSCs in the brain. **(A)** The schematic of mEPSC recording in GL261 in the brain. **(B)** The representative mEPSC recordings in GL261 in **(C-D)** Cumulative distributions of mEPSC inter-event intervals and amplitudes in WT (black, *n* = 16-17 cells in 8 mice) and Asic1a^-/-^ (red, *n*=19 cells in 8 mice) amygdala slices. Insets are the summarized data of inter-event intervals, * p = 0.0205 *(t = 2.431, df = 34)*, and amplitudes, * p = 0.0485 *(t = 2.049, df = 33)*. **(E-F)** The western blot analysis shows the GluA1 **(E)** and GluA4 **(F)** expression levels in GL261 and cultured SY5Y neurons. ꞵ-actin was used as a loading control (same ꞵ-actin bands for both GluA1 and GluA4 results). All statistics were analyzed by a two-tailed unpaired Student’s t-test. Data are mean ± SEM.

### Acid stimulation on neurons induces glioma EPSCs

Drawing from existing references that highlight the role of physiological low pH environments (pH 6-7) in eliciting action potential (AP) spikes in neurons (Vukicevic and Kellenberger, 2004; Du et al., 2014), our study sought to investigate the hypothesis that acid stimulation of neurons can activate glioma postsynaptic activity. We applied direct stimulation of neurons using an acid puff at pH 6.5, which allowed us to monitor and record the ensuing neuronal responses, as visually demonstrated in **Fig. 5A**. The neurons exhibited action potentials in response to the acid puff stimulation, a phenomenon that could be effectively quelled by the application of PcTx1 (200 ng/ml), an Asic1a-specific blocker, and in Asic1a^-/-^ neurons (**Fig. 5B, C**).

**Figure 5.**
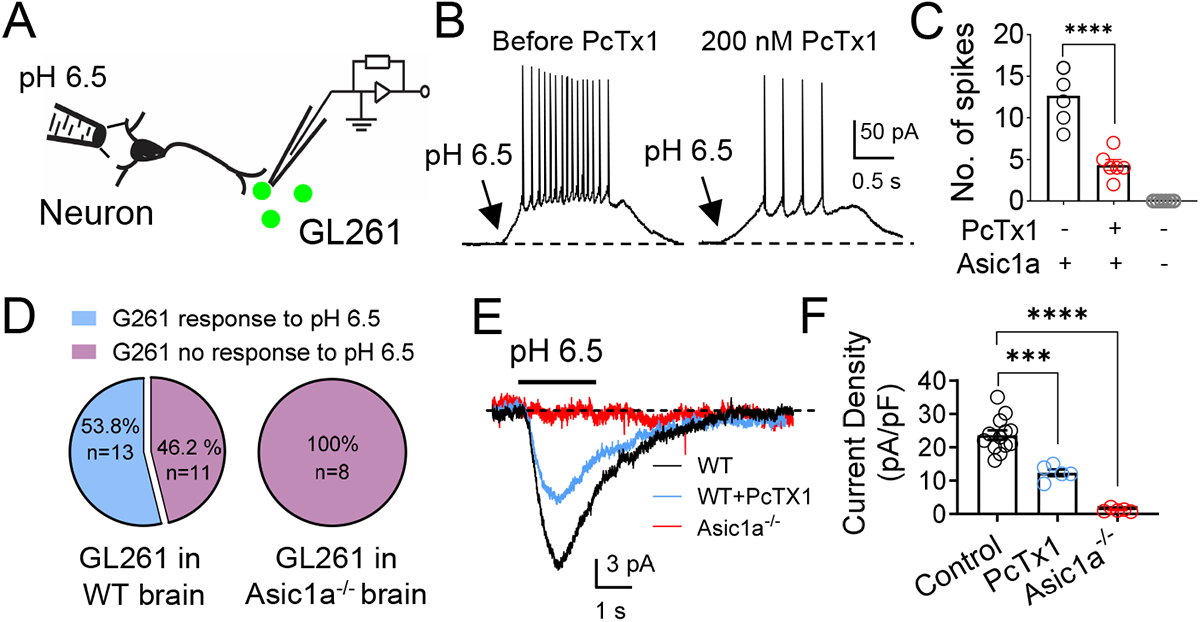
Acid stimulation in neurons induces EPSCs in gliomas in the brain. **(A)** The schematic shows the experimental approach for recording GL261-EPSCs by acid-stimulating neurons (pH 6.5, 2 s). **(B)** The representative results of neuronal action potential induced by acid stimulation with or without 200 nM PcTx1 pre-treatment. **(C)** Summarized data of neuronal action potential spikes in WT and Asic1a^-/-^ neurons with or without PcTX1 (*n* = 5 cells in 3 mice). **** p < 0.0001 *(F (DFn, DFd) 8.448 (2, 15))*. **(D)** The numbers and percentages of GL261 show EPSCs that respond to neuronal acid stimulation in WT and Asic1a^-/-^ brains. **(E-F)** The representative EPSCs that respond to neuronal acid stimulation in WT (black, *n* = 13 cells in 5 mice) and Asic1a^-/-^ (red, n = 5 cells in 5 mice) brains with or without PcTx1 (blue, *n* = 5 cells in 5 mice). *** p = 0.0001, **** p < 0.0001, *(F (DFn, DFd) 3.378 (2, 20))*. All statistics were analyzed by One-way ANOVA Tukey’s multiple comparisons test. Data are mean ± SEM.

Building upon these pivotal findings, we further investigated the neuron-glioma interaction, wherein we introduced pH 6.5 acid puff stimulation on neurons and recorded the EPSCs in the implanted GL261-GFP cells in the amygdala. By stimulating the cortical input neurons using the pH 6.5 puff and recording the EPSC responses of GL261-GFP cells, we identified the effects of acid-induced stimuli via neuron-glioma interactions. In GL261-GFP cells implanted in WT brains, 53.8% of the glioma-EPSC responses were characterized as positive following acid stimulation of neurons, whereas 46.2% exhibited non-responses. In stark contrast to the GL261-WT brains, GL261-GFP cells implanted in Asic1a^-/-^ brains exhibited a complete absence of positive response to the acid-stimulated cortical input (**Fig. 5D-F**). The positive EPSC responses were effectively blocked upon PcTx1 administration (**Fig.5E-F**), further indicating that the acid-mediated responses crucial for glioma-neuron interactions are modulated by Asic1a.

Together, these findings underline the role of low pH environments in orchestrating glioma-neuron interactions and underscore the crucial influence of brain Asic1a in mediating these processes. The intricate connections between pH dynamics, Asic1a, and the activity dynamics of glioma cells are illuminated through this study, thereby laying the groundwork for novel therapeutic avenues aimed at modulating glioma progression.

### *Asic1a* deletion in the mouse brain alters K^+^ channels in the implanted glioma cells

K^+^ channels govern a wide range of biological processes and play critical roles in a variety of diseases by controlling potassium flow across cell membranes (Zhorov, 2011). K^+^ channels are selectively expressed in numerous tumor cells, including gliomas, and impact tumor biological processes, such as proliferation, apoptosis, differentiation, and invasions (Pardo and Stuhmer, 2014; Takayasu et al., 2020; Sakellakis and Chalkias, 2023).

To evaluate the hypothesis that the deletion of Asic1a in neurons inhibits K^+^ channel activity in glioma cells, we conducted a comprehensive analysis focused on voltage-activated K^+^ channels. These experiments involved the recording of voltage-dependent K^+^ currents ranging from -70 to +90 mV in 20 pA increments, within individual GL261-GFP single cells (**Fig. 6A**). Interestingly, GL261-GFP cells implanted in WT brains exhibited significantly larger K^+^ currents in comparison to the K^+^ currents observed in cultured GL261-GFP single cells (**Fig. 6B, C**). Diminished K^+^ current was evident in GL261-GFP cells implanted in Asic1a^-/-^ brains when compared to their GL261-WT counterparts (**Fig. 6B, C**). Although the K^+^ current profiles (current density) between cultured GL261-GFP cells and GL261-GFP cells implanted in Asic1a^-/-^ brains demonstrated statistically significant differences, the discrepancies are smaller than in the GL261-GFP in WT brains. (**Fig. 6B, C**). To further test the role of K^+^ channels, we administered 200 μM 4-aminopyridine (4-AP), a commonly used K^+^ channel blocker (Mitterdorfer and Bean, 2002). which unveiled the presence of distinct K^+^ currents within each experimental group. Specifically, cultured GL261-GFP cells exhibited a combination of delayed A-type (26.67%) and delayed rectifier (73.3%) K^+^ currents (**Fig. 6D, E**). Similarly, GL261-GFP cells implanted in WT brains displayed a combination of 46.43% delayed A-type, 25% delayed rectifier, and 28.57% intermediate rectifier K^+^ currents (**Fig. 6D, F**). Remarkably, the intermediate rectifier K^+^ current was a unique feature solely observed in the glioma-implanted mice model (**Fig. 6E, F**).

**Figure 6.**
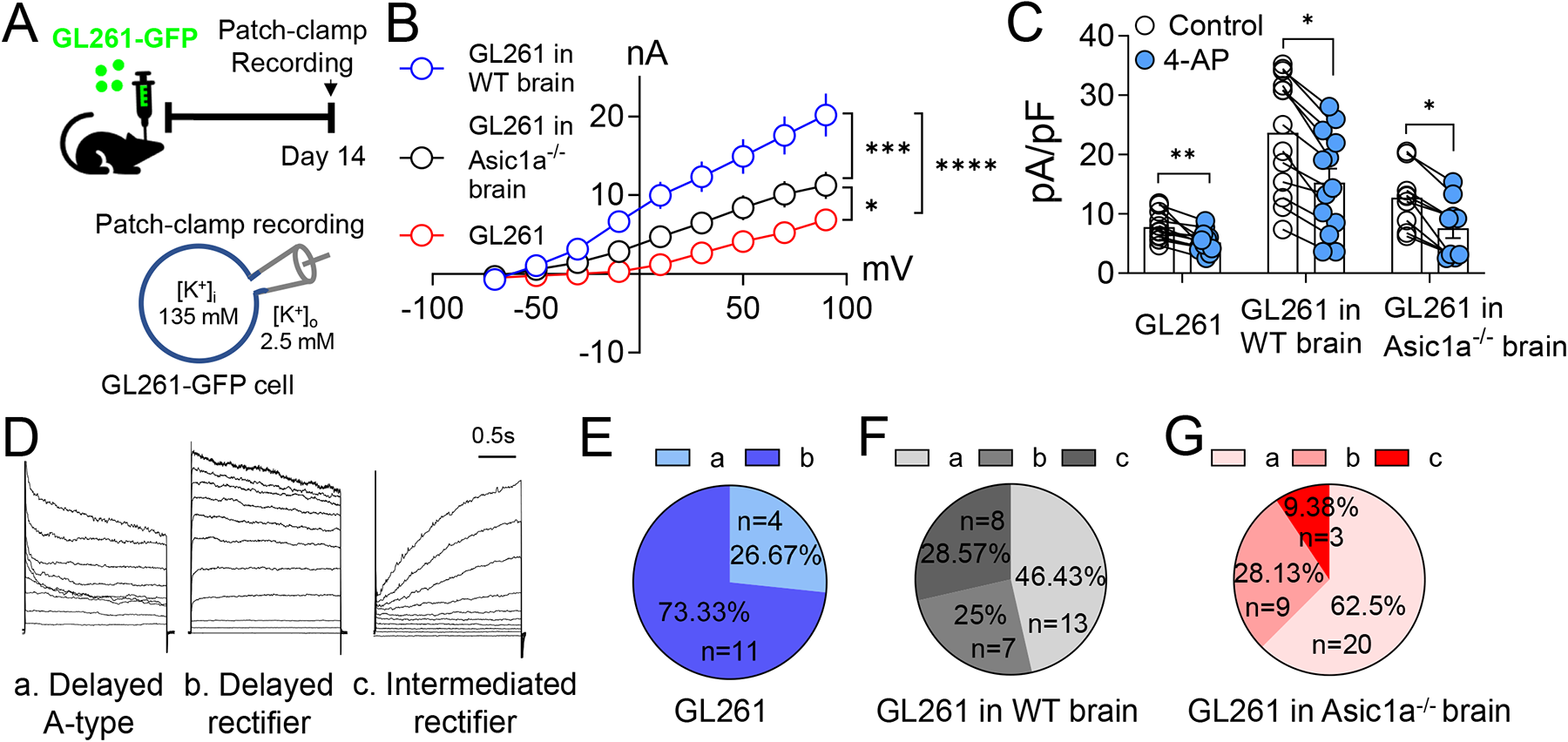
Brain Asic1a deletion reduces K^+^ currents in glioma in the brain. **(A)** The Schematic shows GL261 injection in the brain and patch-clamp recording timeline. **(B)** The I-V curve of K^+^ currents in GL261 cells. Currents were induced by 20 mV membrane potential steps ranging from -70 to +90 mV. Distinct K^+^ currents in cultured GL261 cells (red, *n* = 11 cells), GL261 in WT brains (blue, *n* = 13 cells in 4 mice), and GL261 in Asic1a^-/-^ brains (black, *n* = 9 cells in 4 mice). The current density at +70 mV: cultured GL261 *vs* GL261-WT brain, **** p *<* 0.0001; cultured GL261 *vs* GL261-Asic1a^-/-^ brain * p *=* 0.043; GL261-WT brain *vs* GL261-Asic1a^-/-^ brain *** p *=* 0.0005. The indicated p values represent a two-way ANOVA followed by the Tukey multiple comparisons test. **(C)** The summarized K^+^ current density in these groups with or without 200 uM 4-AP (K^+^ channel blocker). ** p = 0.0045, * p = 0.0248, * p = 0.0440, respectively, by a two-tailed unpaired Student’s t-test. **(D)** The representative types of K^+^ currents were observed in each group. a. Delayed A-type (left), b. delayed rectifier (middle), and c. intermediated rectifier (right). **(E-G)** The Percentage of different types of K^+^ currents observed in cultured GL261 **(E)**, GL261 in WT brains **(F)**, and GL261 in Asic1a^-/-^ brains **(G)**. Data are mean ± SEM.

Further insights arose when comparing GL261-GFP cells implanted in Asic1a^-/-^ brains (62.5% delayed A-type, 28.13% delayed rectifier, and 9.38% intermediate rectifier K^+^ currents) to their GL261-WT counterparts. Specifically, the presence of intermediate rectifier K^+^ currents was significantly reduced in the GL261-Asic1a^-/-^ group relative to the GL261-WT group, 9.38% versus 28.57%, respectively (**Fig. 6F, G**). This finding suggests that the intermediated rectifier K^+^ currents might play a role in influencing glioma-neuron communications, and the effects are Asic1a-dependent.

Together, these findings provided evidence for the decreased K^+^ currents in glioma-implanted Asic1a^-/-^ brains, which could potentially disrupt glioma signaling pathways through neuron-glioma interactions. The presence of distinct K^+^ current profiles, notably the intermediate rectifier currents unique to the glioma-implanted mice, underscores the intricate nature of ion channel dynamics within the context of neuron-glioma interactions.

### *Asic1a* deletion in the mouse brain alters glioma mitogens

The interaction between neurons and glioma cells hinges on the activity-regulated cleavage of neuroligin-3 (NLGN3) that drives glioma progression (Venkatesh et al., 2015; Venkatesh et al., 2017; Pan et al., 2021). This phenomenon has emerged as a crucial mediator, as evidenced by the robust correlation between *NLGN3* expression and survival in glioblastoma patients, underlining its significance in glioma pathophysiology (Derks et al., 2018). In addition to NLGN3, many other molecules, including TGF-β, also function as glioma mitogens (Yang et al., 2022). We observed a reduction in NLGN3 expression within the tumor regions of GL261-GFP implanted Asic1a^-/-^ brains relative to their WT counterparts using immunohistochemistry (**Fig. 7A, B**). This observation was further supported by NLGN3 western blotting data (**Fig. 7C**).

**Figure 7.**
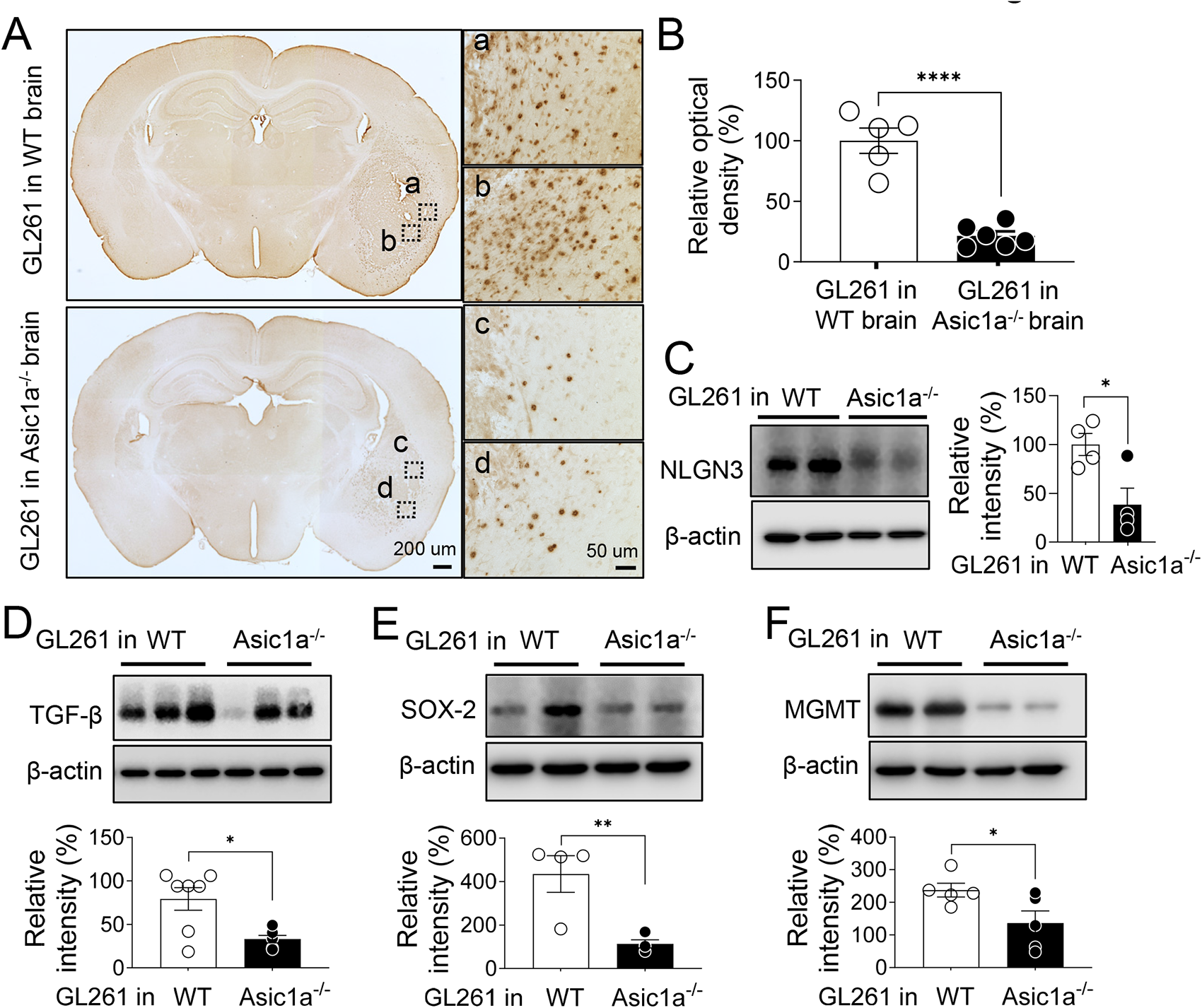
Brain Asic1a deletion alters glioma mitogens. **(A)** The representative immunostaining for neuroligin-3 (NLGN3) in glioma-bearing WT (up) and Asic1a^-/-^ (down) brain slices. **(B)** The relative density of NLGN3 in the amygdala (*n* = 5-6 mice in each group). **** p < 0.0001 *(t=7.578, df=9)*; **(C)** The western blot analysis showing the NLGN3 expression levels in glioma bearing WT and Asic1a^-/-^ brain slices. * p = 0.0232 *(t=3.027, df=6)*. **(D-F)** The western blot analysis shows the TGF-β **(D)**, SOX-2 **(E)**, and MGMT **(F)** expression levels in the glioma-bearing WT and Asic1a^-/-^ brain slices. β-actin was used to demonstrate equal loading in all western blot experiments. The graphs show the quantification of relative signal intensities (*n* = 5-7 mice in each group). All the western blots were cropped for clarity. In D, * p = 0.0158 *(t=3.328, df=6)*; In E, ** p = 0.0099 *(t=3.717, df=6);* In F, * p = 0.0454 *(t=2.368, df=8)*. Data are mean ± SEM.

Broadening the scope of our investigations, we delved into other potential factors that might contribute to high-grade glioma growth, including TGF-β, SOX-2, and O^6^-methylguanine-DNA methyltransferase (MGMT). The TGF-β pathway, known for its multifaceted influence on glioma initiation and progression, was examined due to its impact on diverse aspects like cell proliferation, tumor invasion, angiogenesis, and stemness maintenance in glioma stem cells (Yamada et al., 1995; Kaminska et al., 2013; Joseph et al., 2022). The expression of SOX2, linked to tumor formation, chemoresistance, and tumor stem cell-like characteristics, was also explored (Berezovsky et al., 2014; Boumahdi et al., 2014). Methylation of the MGMT gene promoter has been observed in approximately 50% of glioblastoma, it thus can be used as a biomarker (Butler et al., 2020). Our analyses revealed a reduction in the expression levels of TGF-β, SOX-2, and MGMT within the tumor regions of Asic1a^-/-^ brains compared to their WT counterparts (**Fig. 7D-F**). Collectively, these findings illuminate the intricate dynamics of neuron-glioma interactions, highlighting the extensive regulatory influence that neuronal cells exert on the growth and pathophysiology of glioma.

## Discussion

High-grade gliomas remain one of the most aggressive and challenging brain tumors to treat, requiring novel approaches to improve therapeutic outcomes. The integration of glioma cells into the neuronal microenvironment has been recognized as a significant contributor to tumor progression. Our current study focuses on the neuronal H^+^-Asic1a axis and provides a deeper understanding of how glioma cells exploit this microenvironment to drive its malignant behavior. In this study, we have unveiled insights into the role of neuronal H^+^ signaling pathways in glioma progression and its intricate interaction with neural elements. By elucidating the molecular mechanisms involved in the formation and function of neuron-to-glioma synapses, the study enhances our understanding of how glioma cells actively interact with and respond to their surroundings.

### Glioma growth regulates the neural pH microenvironment

As the tumor expands, it creates a unique microenvironment characterized by regions of low pH, primarily due to the Warburg effect -a phenomenon where cancer cells metabolize glucose predominantly through glycolysis, even in the presence of oxygen (Liberti and Locasale, 2016). This metabolic shift results in the accumulation of lactic acid, leading to an acidic environment around the tumor cells. Consolidated information from several studies has brought forth the knowledge that tumor masses can demonstrate heterogeneous low pH levels ranging from 7.4 to 6.2 units, in general, that can fall as low as 5.5-3.4 units and even below (Webb et al., 2011; Kato et al., 2013; Chen et al., 2014). Interestingly, this acidic microenvironment has been observed to exert a paradoxical effect on surrounding neurons. While low pH is generally detrimental to neuronal function, studies suggest that in proximity to glioma growth, certain neurons can undergo adaptive changes. These neurons display enhanced excitability and synaptic transmission in response to the acidic environment (Du et al., 2014). The underlying mechanisms driving this phenomenon are multifaceted. It is hypothesized that neurons in the vicinity of glioma cells may upregulate specific ion channels and receptors, such as ASICs, as a response to the low pH, thereby modulating their activity to sustain communication within the affected neural circuitry. Our data support this concept, we found that brain Asic1a expression and activity are elevated following glioma implantation. This adaptation might be an attempt to counteract the inhibitory effects of the acidic environment on neuronal signaling. Understanding how glioma-induced low pH impacts neuronal function is a complex area of ongoing research. ASICs are also expressed in glioma and can regulate tumor progression, whereas controversial pro- and anti-cancer effects were reported (Honasoge and Sontheimer, 2013; King et al., 2021; Sheng et al., 2021). Alternative mechanisms are needed to explain how glioma acidity modulates its pathogenesis. The advantage of our system is that we only knock out Asic1a in the microenvironment of neurons while keeping Asic1a intact in glioma cells so that we can examine how brain Asic1a affects glioma behaviors. In the future, it may be interesting to knock out Asic1a in both neurons and glioma cells to see whether this double Asic1a deletion can facilitate the effects of H^+^ on glioma progression.

### Role of brain Asic1a in glioma progression

The current study provides compelling support for evidence that neuron-to-glioma synapses drive glioma progression. Previous research has established that glutamatergic synapses can influence tumor growth and invasion, yet the molecular mechanisms governing these interactions are not fully understood. We identified brain Asic1a as a key mediator in this process, which sheds new light on the intricate interplay between neuronal activity and high-grade glioma malignancy. The observation that brain *Asic1a* deletion leads to increased survival in glioma-bearing mice underscores the complex interaction between tumor biology and neural circuitry. Despite the absence of discernible differences in body weight and brain weight at day 14 post-glioma implantation, significant differences emerged in later stages (**Fig. 1**). The reduction in body weight specifically between days 30 and 36 in the glioma-bearing WT mice could indicate late-stage manifestations of the tumor’s systemic effects. Conversely, the significantly higher survival rate observed in Asic1a^-/-^ mice with glioma implantation indicates the role of brain Asic1a in influencing growth and progression. The decrease in glioma cell size, observed through Nissl staining and reduced GFP expression, provides structural evidence for the impact of brain Asic1a deletion on glioma growth (**Fig. 1 and 2**). These changes might indicate altered cellular processes such as proliferation and invasion. Further investigations are needed to elucidate the underlying mechanisms by which the host’s Asic1a influences glioma growth and the potential contribution of Asic1a to glioma cell signaling pathways. A particularly intriguing aspect of this study is the observed role of brain Asic1a in modulating voltage-activated K^+^ channels in the glioma microenvironment (**Fig. 6**). Numerous studies have shown that pharmacological block of voltage-gated K*^+^* channels reduces the proliferation of cancer cells (Fraser et al., 2000; Abdul and Hoosein, 2002; Ouadid-Ahidouch and Ahidouch, 2008; Menendez et al., 2010; Shah et al., 2021; Sheng et al., 2021). The decrease in K^+^ current density in glioma cells in Asic1a^-/-^ mice hints at Asic1a’s potential influence on glioma channel functions. The presence of distinct K^+^ current types in both glioma-implanted mice groups highlights the intricate regulatory mechanisms underlying K^+^ currents in this context. Its decrease in glioma cells implanted in Asic1a^-/-^ mice suggests a possible role for this K^+^ current under the communication between glioma and neurons. Further investigations into the functional properties and mechanisms of these K^+^ currents could offer insights into how Asic1a affects glioma-neuron crosstalk and the overall tumor behavior.

### Role of brain Asic1a in neuron-to-glioma synapses

The molecular mechanisms underlying the modulation of neuron-to-glioma synapses by Asic1a raise intriguing questions. How does Asic1a activation influence the release of neurotransmitters or other signaling molecules from neurons to affect glioma cells? Are there downstream signaling pathways that are activated upon Asic1a stimulation, and do these pathways contribute to the observed changes in glioma progression? Identification of these pathways could provide valuable insights into the broader molecular context of glioma-neuron interactions.

Neuron-glioma synapses are related to both chemical and electrophysiological responses. Asic1a activation has been demonstrated to modulate synaptic neurotransmitter release in response to extracellular acidification (Storozhuk et al., 2016; Storozhuk et al., 2021). This modulation occurs through several mechanisms, including the regulation of presynaptic Ca^2+^ levels and the release probability of synaptic vesicles (Sudhof, 2013). Changes in synaptic neurotransmitter release can have downstream effects on synaptic plasticity and the organization of synapses (Citri and Malenka, 2008). If Asic1a activation leads to alterations in presynaptic neurotransmitter release, it could potentially influence the signaling and interaction between pre- and postsynaptic proteins, including NLGN3, which is a postsynaptic cell adhesion molecule that plays a role in synapse formation and function and promotes the expression of synapses related gene in glioma cells (Venkatesh et al., 2015). Consistent with these findings, we found that there is decreased NLGN3 expression in glioma-bearing Asic1a^-/-^ mice (**Fig. 7**). Neuronal hyperactivity promotes glioma proliferation by activating ion channels through glutamate secretion (Hua et al., 2022). These hyperexcitable neurons induce evoked ESCs in glioma cells mediated by AMPAR (Venkataramani et al., 2019; Venkatesh et al., 2019). The average AMPAR current density in glioma cells was reduced in the glioma-bearing Asic1a^-/-^ mice compared to the WT mice. Similarly, we tested whether functional neuron-glioma synapses exist when we implanted glioma cells into the amygdala. We found that an evoked-AMPAR current exists at the leading edge of the tumor.

Our study also reveals the influence of Asic1a on glioma AMPAR currents. Glutamate, particularly through AMPAR, is known to play a pivotal role in the crosstalk between glioma cells and neurons (Venkataramani et al., 2019; Venkatesh et al., 2019). The reduction in evoked-AMPA current density and mEPSC frequency in glioma cells implanted in the Asic1a^-/-^ mice, suggests that Asic1a deletion hampers efficient communication between glioma cells and neurons. These findings underscore the intricate balance maintained by Asic1a in facilitating proper neurotransmission in the tumor microenvironment. The preservation of mEPSC amplitude suggests that post-synaptic mechanisms remain relatively unaffected by Asic1a deletion, emphasizing the specificity of its impact on neural communication.

### Therapeutic implications and future directions

Our finding that pharmacological targeting of brain Asic1a inhibits glioma progression holds promising therapeutic implications. Developing drugs that modulate Asic1a activity could represent a novel avenue for high-grade treatment by disrupting the neuron-to-glioma signaling axis. However, further investigation is needed to better understand the specific effects of Asic1a modulation on neuronal activity, synaptic transmission, and overall CNS function.

### Limitations and future challenges

While the study presents a groundbreaking advancement in the field, certain limitations warrant consideration. The current findings provide insights into the role of Asic1a in neuron-to-glioma communication, but the complexity of synapse formation, synaptic plasticity, and the involvement of other ion channels and receptors cannot be overlooked. Additionally, the study’s in vivo observations should be interpreted with caution, as the tumor microenvironment is inherently heterogeneous and subject to multiple variables.

## Conclusion

In conclusion, this study elucidates the multifaceted role of Asic1a in high-grade glioma progression and its interaction with neural elements. The increased survival rates, alterations in K^+^ channel activity, disruptions in AMPA receptor currents, and responses to acidic microenvironments collectively underscore Asic1a’s multifunctional involvement in glioma pathophysiology. These findings hold promising implications for the development of novel therapeutic strategies targeting Asic1a to mitigate glioma progression. Further mechanistic investigations are warranted to elucidate the precise molecular mechanisms driving these effects and to assess the translational potential of Asic1a-targeted interventions in the clinical setting. Collectively, our study paves the way for future research aimed at harnessing the potential of Asic1a manipulation for improving glioma management and patient outcomes.

## AUTHOR CONTRIBUTIONS

J.D. and Y.P. conceived the project. J.D., Y.P., G.P., Q.G., and Z.J. designed the experiments. G.P. and J.D. performed the patch-clamp experiments and data analysis. Z.J. and Q.G. performed the behavior experiments and data analysis G.P., J.D., and Y.P. wrote the manuscript. All authors reviewed and edited the manuscript. Method to determine the equal contribution authorships: G.P. finished figures 2-6. Z.J. finished figures 1 and 7 and all the mouse surgeries.

## ACKNOWLEDGMENTS

We thank Dr. John Wemmie at the University of Iowa for providing the ASIC1a^f/f^ mice. We thank Drs. Elizabeth Tolley and Rongshun Zhu in BERD Clinic at UTHSC for statistical support. J.Du. is supported by the National Institutes of Mental Health (1R01MH113986), the Cystic Fibrosis Foundation (002544I221), and the University of Tennessee Health Science Center start-up fund.

